# Csde1 cooperates with Strap to control translation of erythroid transcripts

**DOI:** 10.1101/203539

**Authors:** Kat S. Moore, Nurcan Yagci, Floris van Alphen, Alexander B. Meijer, Peter A.C. ‘t Hoen, Marieke von Lindern

## Abstract

Erythropoiesis is regulated at many levels, including control of mRNA translation. Changing environmental conditions, such as hypoxia, or the availability of nutrients and growth factors, require a rapid response enacted by the enhanced or repressed translation of existing transcripts. Csde1 is an RNA-binding protein required for erythropoiesis and strongly upregulated in erythroblasts relative to other hematopoietic progenitors. The aim of this study is to identify the Csde1-containing protein complexes, and investigate their role in regulating the translation of Csde1-bound transcripts. We show that Strap, also called Unrip, was the protein most strongly associated with Csde1 in erythroblasts. Strap is a WD40 protein involved in signaling and RNA splicing, but its role is unknown when associated with Csde1. Reduced expression of Strap did not alter the pool of transcripts bound by Csde1. Instead, it reduced the mRNA and/or protein expression of several Csde1-bound transcript, that encode for proteins essential for translational regulation during hypoxia, such as Hmbs, eIF4g3 and Pabpc4. Also affected by Strap knockdown were Vim, a Gata-1 target crucial for erythrocyte enucleation, and Elavl1, which stabilizes *Gata-1* mRNA. Thus, we found that the Csde1/Strap complex is at the crossroad of multiple pathways governing translation in erythroblasts.

## Introduction

Maintenance of correct numbers of erythrocytes in peripheral blood requires continuous replenishment with newly synthesized cells. Proliferation and differentiation of erythroblasts needs to be tightly balanced to prevent anemia and ischemic damage of organs, or an excess of erythrocytes and a risk for stroke. Environmental factors such as growth factors (e.g. erythropoietin and stem cell factor) or nutrients (e.g. iron) are crucial to control erythropoiesis, which occurs in part through control of translation of the available transcriptome. RNA binding factors have an important role in control of translation. For instance, iron regulatory proteins 1 and -2 (Irp1, Irp2) bind to the iron response element in *Ferritin* and *Transferrin receptor* mRNA to control expression of the encoded proteins that are crucial to erythropoiesis [1]. Zinc finger binding proteins 36 like 1 and -2 (Zfp36l1, Zfp36l2) bind to a large number of transcripts and deletion of Zfp36l2 disrupts erythropoiesis [2,3]. The RNA-binding protein Csde1 (cold shock domain protein e1), first described as Unr (upstream of Nras) [4], is widely expressed, but expression levels differ per cell type. In the hematopoietic system, expression of Csde1 is increased more than 100-fold in erythroblasts relative to other hematopoietic cells, and expression of Csde1 is reduced in Diamond Blackfan Anemia [5]. Knockdown of Csde1 impairs both proliferation and differentiation of erythroblasts [5].

Csde1 regulates the fate of target transcripts by binding to the 3′ UTR [4,6,7] or to IRESs (Internal Ribosomal Entry Sites) [8–10]. Because Csde1 is capable of binding a broad variety of mRNAs containing A/G-rich binding motifs, it is likely that it functions as a global regulator of translation [11,12], By consequence, it is involved in diverse processes including sex determination in Drosophila [13], cell cycle control [10], and control of metastasis in melanoma [14]. We recently identified the transcripts bound by Csde1 in erythroblasts. These transcripts encoded proteins involved in protein homeostasis: translation factors, ribosome biogenesis factors, subunits of the proteasome and peptidases [15].

The function of RNA-binding proteins depends on associated proteins. For instance, RNA binding proteins that interact with AU-rich elements in the 3′UTR of transcripts such as AUF1 (AU-rich element binding factor) can interact either with the pre-initiation scanning complex to enhance translation, or with the Cnot1 (Ccr4/Not complex 1) complex which results in deadenylation [16]. Similarly, the role of Csde1 is likely influenced by associated proteins that may affect its RNA-binding affinity and/or functional consequences. Csde1 cooperates with Pabp (PolyA binding protein) when interacting with the 3′ UTR [17,18] and with PTB (polypyrimidine tract binding protein) and hnRNP (heterogeneous nuclear binding protein) C1/C2 when interacting with internal ribosomal entry sites (IRESs) [7,10,19]. It also interacts with Strap (serine-threonine receptor associated-protein, also called Unrip) [20]. Strap is a member of a large family of WD40 (Trp/Asp) repeat containing proteins that are known to function as relatively promiscuous adapters. WD40 domains function as a platform for protein/protein interactions. The association of WD40 domains with phosphorylated Ser-Thr residues often place WD40 domain proteins in network nodes of signaling cascades [21]. Strap is involved in numerous biological pathways, including TGFβ signaling [22,23], MAPK signaling [24], Wnt signaling [25], Notch signaling [26] and assembly of the survival motor neuron (SMN) complex [27]. Its association with the SMN complex is mutually exclusive with Csde1 binding. Strap also associates with 4E-T (eIF4E Transporter) together with Csde1 to promote P-body assembly [28]. Apart from P-body assembly, little is known about the function of Strap when bound to Csde1.

The aim of this study was to investigate Csde1 protein complex formation in erythroblasts. Strap was the most strongly associated protein in mouse erythroblasts. Strap knockdown did not influence Csde1 target transcript binding, but reduced protein expression of many Csde1-bound transcripts and enhanced expression of some other targets. Proteins whose expression was regulated by Strap are involved in terminal erythroid differentiation, cell cycle regulation, and the hypoxic response.

## Results

### Strap strongly associates with Csde1

To investigate which proteins form a complex with Csde1 in erythroblasts, we utilized the strong affinity of streptavidin-biotin interaction as an efficient alternative to antibody-based immunoprecipitation to purify Csde1 protein/mRNA complexes [29,30]. BirA and biotagged Csde1 were coexpressed in Mel cells [5]. Csde1-bound protein complexes were enriched on streptavidin-coated Dynabeads under conditions that preserve binding of target mRNAs [15]. Mel cells expressing the biotin ligase BirA without biotagged Csde1 were used as a control.

SDS-PAGE of cell lysates and subsequent silver staining showed a series of bands representing endogenously biotinylated proteins that are common to both the Csde1-pulldown lane and the BirA pulldown control (Fig. 1A). Selectively present in the Csde1-pulldown lane were two bands at ^~^100kDa and ^~^45kDa, representing Csde1 itself and an unknown associated protein, respectively. Mass spectrometry analysis of the bio-Csde1 pulldown eluate and the BirA pulldown control eluate identified proteins enriched in the Csde1 fraction. A two-way t-test, applying an artificial within groups variance (SO) [31] of 0.8 was used to set the threshold for significant binding (Fig. 1B, Table 1). These proteins included Znfx1 (Zinc finger NFx1-type containing 1), Pabpc1 and Pabpc4. Notable is the high fold change enrichment of Strap (Fig. 1C). Strap was nearly twice as abundantly associated with Csde1 in erythroblasts than the next highest enriched protein (Znfx1, predicted mass 220kDa). Strap was previously shown to associate with Csde1/Unr [20], but little is known about how the two proteins cooperate to affect Csde1’s function.

**Fig 1.**
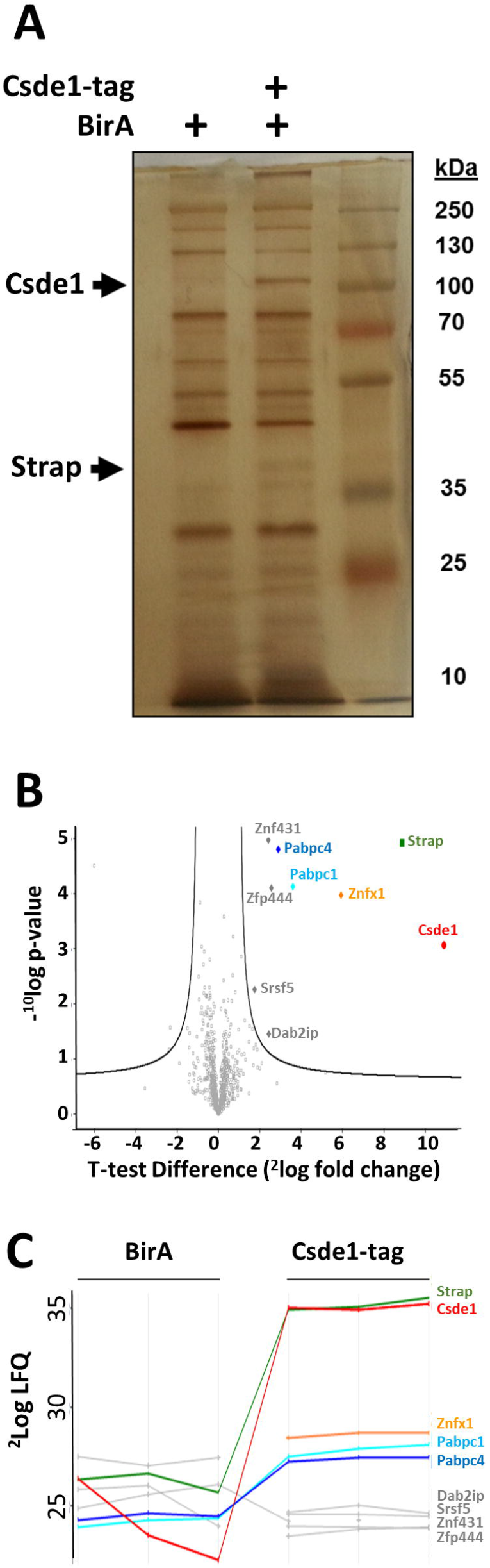
Proteins associated with Csde1 upon pull down of biotagged Csde1. **(A)** Lysate of MEL cells expressing the biotin ligase BirA with or without biotagged Csde1 was incubated with streptavidin beads, washed, and eluted in laemmli buffer before loading on SDS-PAGE and silver staining. Third lane is a protein size ladder. **(B)** Mass spectrometry of proteins pulled down by Csde1, and analysis by two-way t-test revealed 8 Csde1-associated proteins at a significance threshold of S0=0.8. −10log p-value is plotted against 2log fold change (n=3).**(C)** Protein profile expression plot of Csde1-associated proteins (LFQ: Label free quantification). Lines are discontinued when peptides were not detected in control pull down in BirA only MEL cells

**Table 1:**
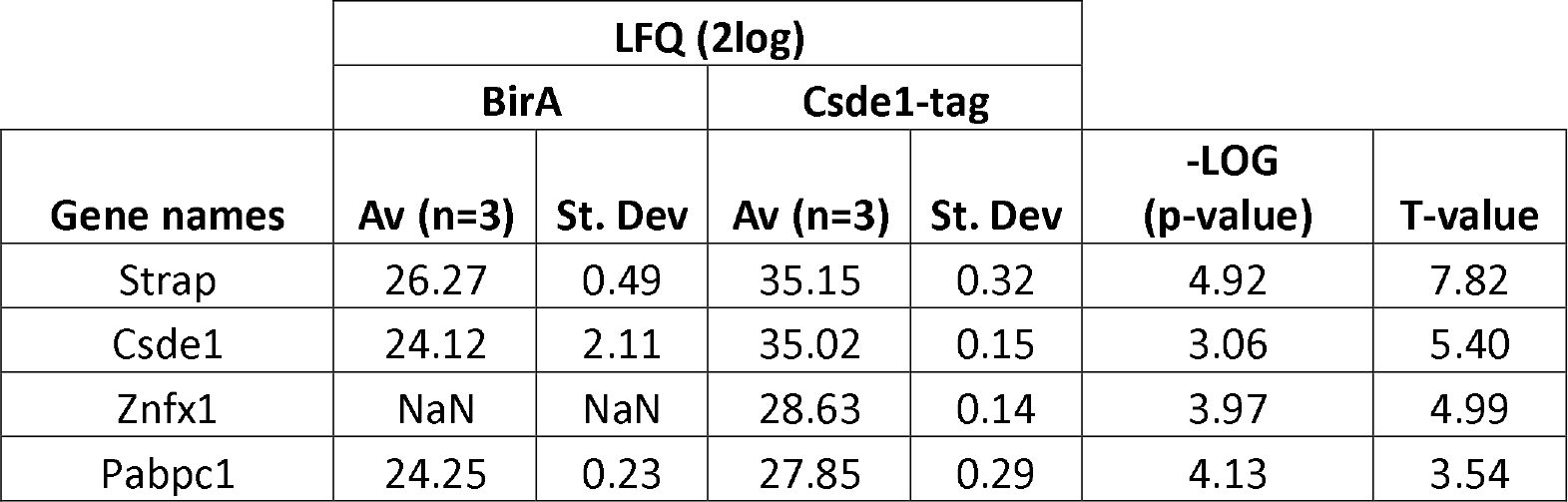
Significant proteins from MS Csde1 pulldown vs BirA

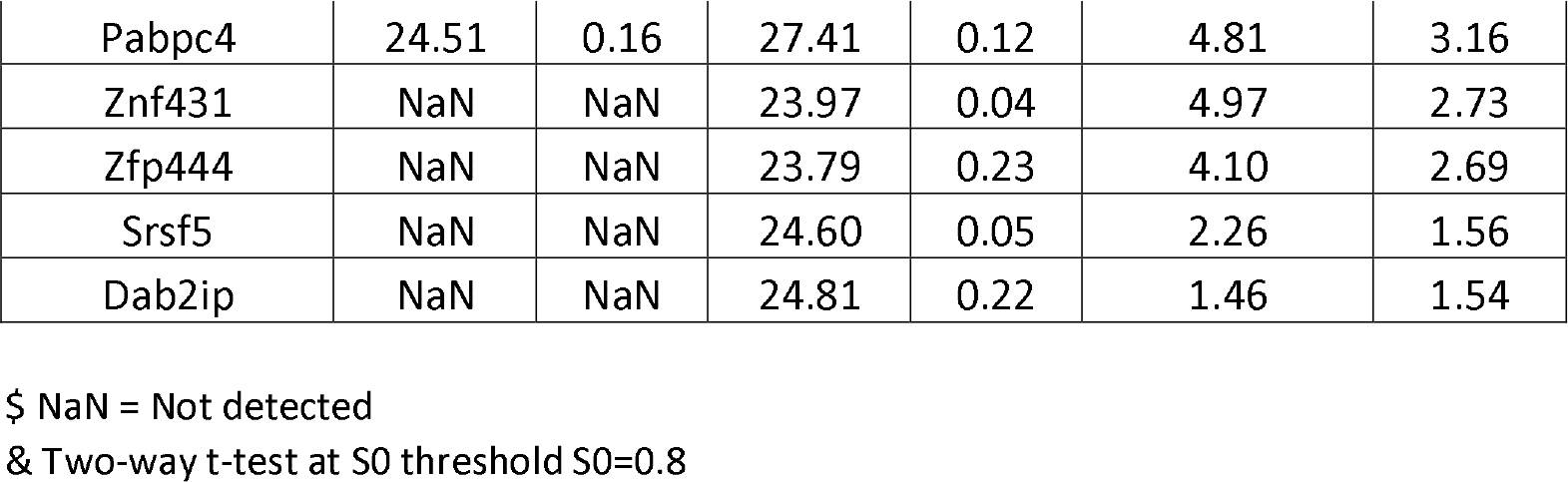

### Csde1 does not sequester Strap from the nucleus

The association of Strap with Csde1 is exclusive with binding to survival motor neuron (SMN) complex protein Gemin7 [27], suggesting that Csde1 may compete with other pathways for binding to Strap. We investigated whether Strap localization is influenced by the abundance and functionality of Csde1. Strap was co-precipitated with Csde1 (Fig. 2A). Whereas Csde1 was largely depleted from the supernatant following pull down, Strap partially remained in the supernatant, suggesting that Csde1 does not sequester the total pool of cytoplasmic Strap.

**Fig 2.**
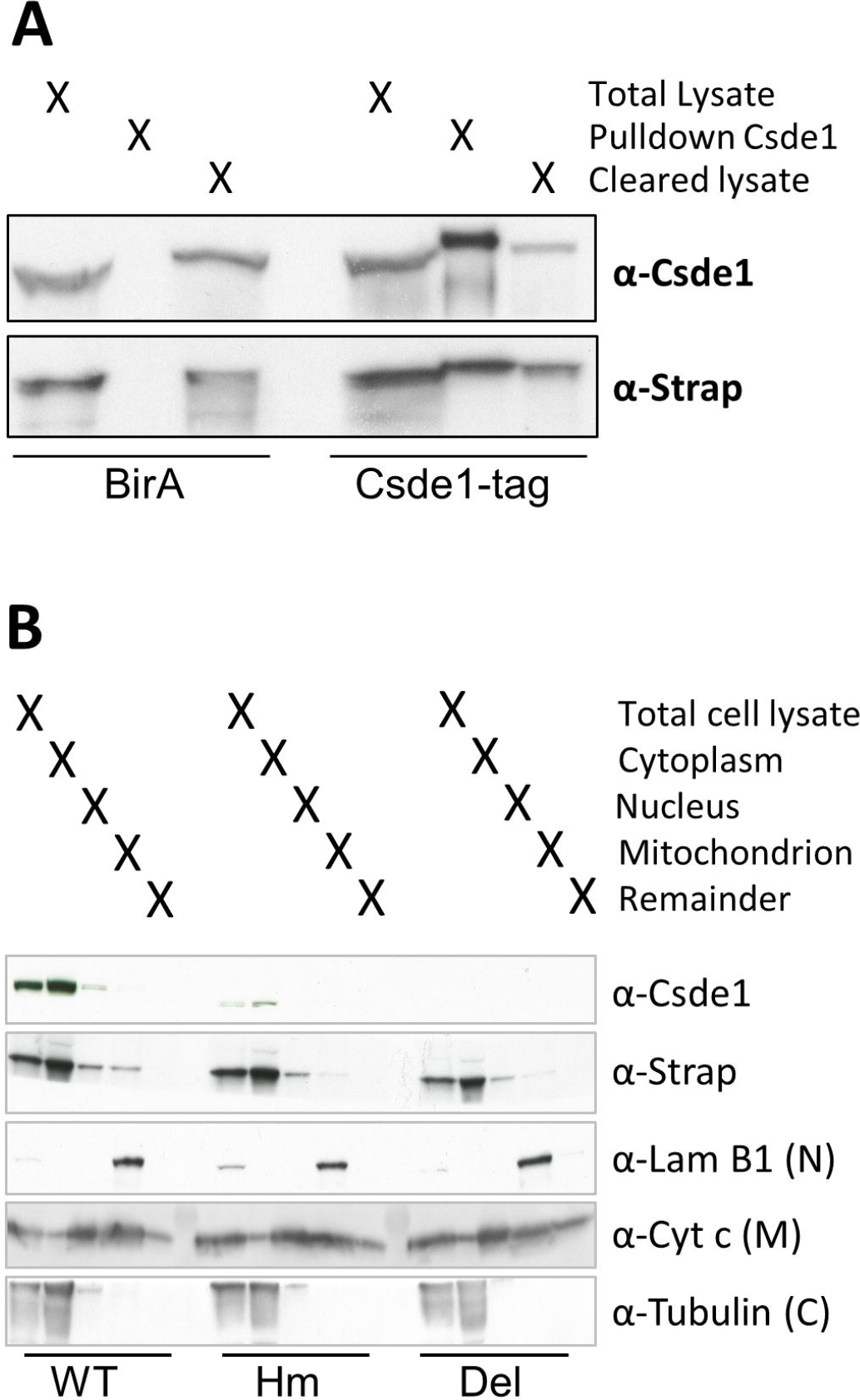
Reduction of Csde1 expression does not alter Strap localization. **(A)** Total cell lysates (**TL**) of MEL cells expressing BirA plus or minus biotagged Csde1 was used to pull down Csde1 using streptavidin beads (**PD**). TL, the PD and the cleared lysate (**CL**) were loaded on SDS-PAGE. Western blots were probed with anti-Csde1 and anti-Strap antibodies. (**B**) Western blot loaded with lysate fractions from parental MEL cells (**WT**), or CRISPR clones with deletions in Csde1 indicated as hypomorphic (**Hm**, in-frame deletion of the 1^st^ cold shock domain), or deleted (**Del**, out-of-frame deletion of the 1^st^ cold shock domain, unexpectedly resulting in low expression of a N-terminally truncated protein). Lysates (T, total lysate) were fractionated into cytoplasmic C), mitochondrial (M), and nuclear (N) extracts. Remaining materials (R) is also loaded. Antibody staining was performed for Csde1, Strap, Lamin B1 (nuclear control), Cytochrome C (mitochondrial control), and Tubulin (cytoplasmic control).

To investigate whether Csde1 expression could alter the distribution of Strap, we also tested subcellular distribution of Strap in MEL cells with a Crispr/Cas-induced in-frame deletion (Hm) in Csde1, in which Csde1 expression was less than 50% of wt MEL levels, or with an out-of-frame deletion (Del) (Fig2B) [15]. MEL cells were fractionated to produce cytoplasmic, nuclear and membrane lysates that were subjected to Western blotting for Csde1, Strap, Lamin B1(nuclear control), Tubulin (cytoplasmic control), and Cytochrome C (mitochondrial control) (Fig. 2B). In wild-type MEL cells, Strap was abundantly localized in the cytoplasm and expressed at lower levels in the nucleus. Csde1 is mainly localized in the cytoplasm and present in small quantities in the mitochondrion. The nuclear fraction of Strap is not increased in Csde1 mutant clones, indicating that Csde1 does not sequester Strap from the nucleus (Fig. 2B).

### Loss of Strap does not affect Csde1’s ability to bind transcripts

We previously identified 292 transcripts that were enriched upon pull down of biotagged Csde1 from MEL cells with a Benjamini-Hochberg false discovery rate (FDR) cutoff of 5% [15]. To investigate whether association with Strap is required for the binding of transcripts by Csde1, Strap expression was reduced in BirA and BirA/biotag-Csde1 expressing MEL cells using a Strap targeted shRNA expressed transduction and a non-targeting control short hairpin (Sc). Knockdown of Strap was confirmed by Western blotting (Fig. 3A). Next, we pulled down biotagged Csde1 from MEL cells expressing BirA only, or BirA plus biotagged Csde1 to identify Csde1 protein complexes in presence and absence of Strap. RNA was isolated from the Csde1 protein complexes and subjected to high-throughput RNA sequencing. Principal component analysis (Fig. 3B) on the 292 previously determined Csde1 targets [15] showed that the presence of biotagged Csde1 in presence or absence of Strap counts for the majority of variance (PC1, 58%), representing the effect of the pulldown versus the control (BirA). In PC2 (21%), Csde1 pulldown samples clustered by replicate, not by the abundance of Strap. Overall, the PCA indicated that Strap knockdown had a weak effect on Csde1 target binding.

**Fig 3.**
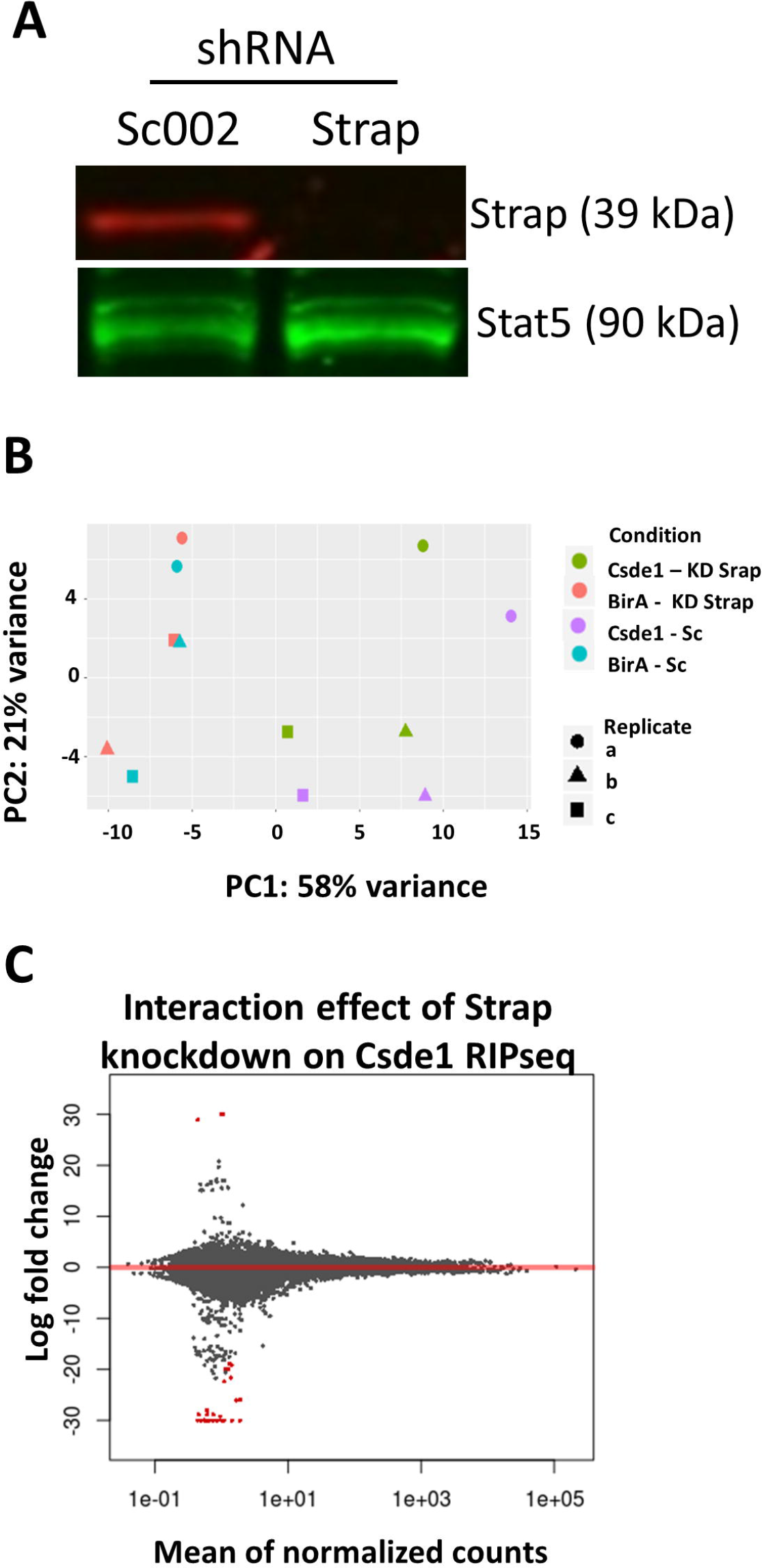
Knockdown of Strap does not affect the pool of Csde1-bound transcripts. **(A)** MEL cells expressing BirA and biotagged Csde1 were treated with a short hairpin targeting Strap or a non-targeting short hairpin (Sc). Knockdown of Strap was confirmed by Western blot staining for Strap and Stat5 as a loading control. **(B)** Principal component analysis of transcripts bound to streptavidin beads incubated with MEL cells expressing BirA with or without biotagged-Csde1, and treated with anti-Strap shRNA (KD Strap) or control shRNA (Sc). Three biologically independent experiments were performed (a-c) **(C)** MA plot of the interacting effect of biotagged Csde1 and anti-Strap shRNA on transcripts pulled down by Csde1. The 2log fold change was calculated from average read counts in presence and absence of Strap (cpm, see B) and plotted against 10log average read counts per million (cpm). Highlighted in red are interaction-term significant transcripts.

To identify Csde1-associated transcripts a Wald test was performed on samples containing biotagged Csde1 versus those without biotagged Csde1 in cells with reduced or control expression of Strap, 183 enriched transcripts were identified as significant (FDR < 0.05) (**S1 Table**). These transcripts were present in relatively high abundance, averaging approximately 2311 cpm across all samples (**S1A Fig**). Only 35 transcripts differed between samples expressing anti-Strap shRNA versus the control shRNA (Sc) (FDR < 0.05) (**S2 Table, S1B Fig**). These transcripts are predominantly expressed at a low abundance (<10 reads) and may not represent biological relevance. To determine whether Strap expression significantly alters the binding of some transcripts by Csde1, we used an interaction model for differential pull down of transcripts by biotagged Csde1 in presence and absence of Strap. The interaction model identified 31 significant transcripts at an FDR cutoff of 5% (Fig. 3C, **S3 Table**). These transcripts were exclusively present in very low abundance across all samples. Importantly, none of the transcripts identified in the interaction model were Csde1 targets (they were not selectively precipitated from lysates containing biotagged Csde1), indicating that Strap does not affect Csde1 mRNA binding.

### Strap controls transcript and protein expression of select Csde1 targets

Strap may affect Csde1 function with respect to mRNA stability and translation. We performed RNA sequencing on total mRNA, and mass spectrometry on cell lysates, after targeting Strap for shRNA-mediated knockdown in MEL cells, using a scrambled shRNA (Sc) to control for the effects of viral exposure. Principal component analysis of RNA sequencing results showed that the majority of variation (87%) can be explained by the knockdown of Strap (**S2 Fig**). After Strap knockdown, 3,828 transcripts were differentially expressed with a FDR < 0.05, of which 70 represented Csde1-associated transcripts (**S4 Table**). Twenty-eight Csde1-bound transcripts increased in expression while 42 Csde1-bound transcripts decreased in expression in response to Strap knockdown. The observed change in expression was not related to the likelihood of being a Csde1 bound transcript (Fig. 4A). Csde1 targets altered by Strap knockdown included transcripts encoding several translation factors (*Eif1*, *Eif2B1*, and *Eif3H*, *Pabpc1*, *Pabpc4*), proteasome subunits (*Psme1*, *Psmc1*, *Psma2*), RNA processing/transport (*Exosc1/Exosc10*, *Thoc5*) and proteins involved in cell cycle control (*Fbxo5*, *Spc24*, *Kif23*), often via (de)ubiquitinylation (*Actr8*, *Ube2c*, *Cdc23*). Also notable were *DEAD-domain protein X18* (*Ddx18*), *Ran-binding protein 1 (Ranbp1)*, *hydroxymethylbilane synthase (Hmbs*, involved in heme biosynthesis), *Platelet factor 4 (Pf4)* and *osteoclast stimulating factor 1 (Ostf1)* (**S5 Table**).

**Fig 4.**
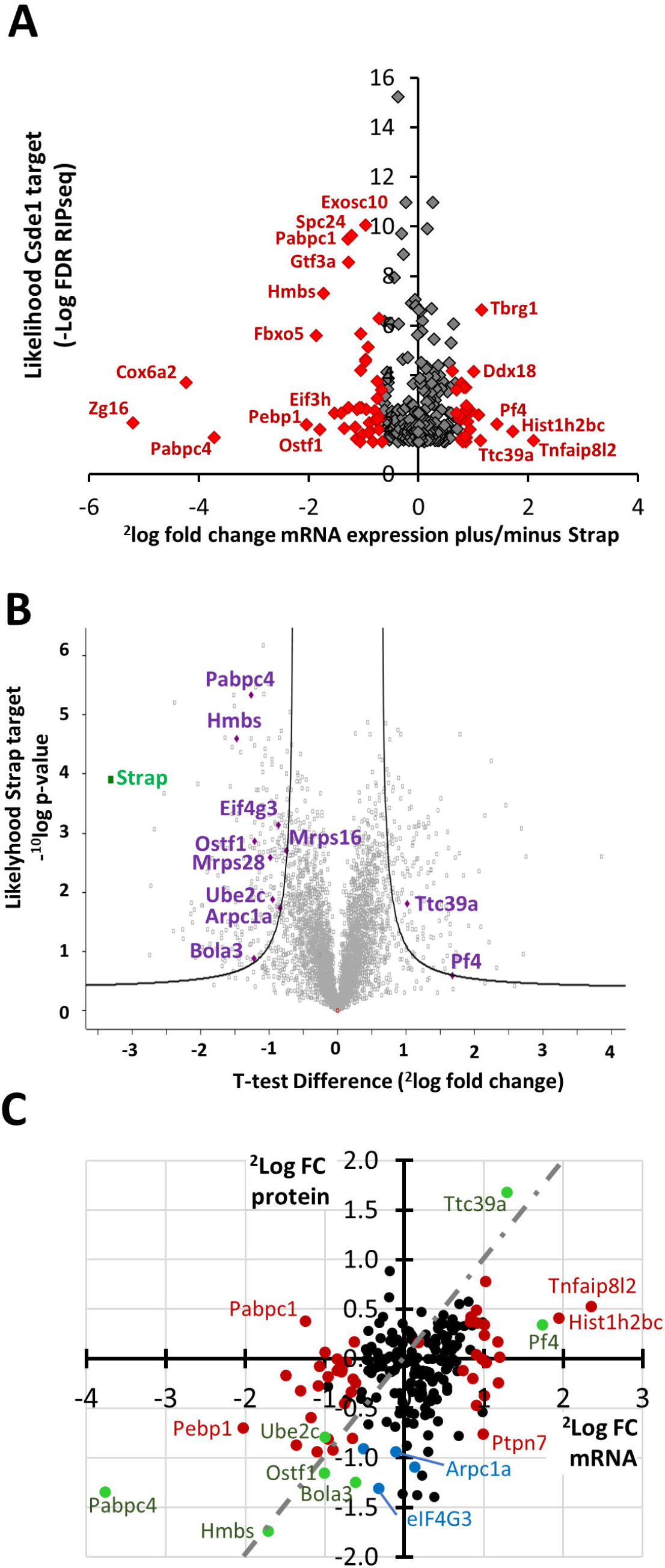
Strap knockdown affects the expression of select Csde1-bound transcripts. MEL cells were treated with shRNA directed against Strap in 3 independent experiments. Cells were processed for transcriptome and proteome analysis by RNA sequencing and mass spectrometry. **(A)** The fold change expression of transcripts in Strap shRNA versus Sc002 shRNA treated MEL cells (2log values; X-axes) was compared to the likelihood of that transcript being bound to Csde1 (−10log values; Y axes), with a cut off at FDR<0.05. Transcripts highlighted in red are also differentially expressed at FDR<0.05. **(B)** The fold change expression of proteins in Strap shRNA versus Sc002 shRNA treated MEL cells (2log values; X-axes) was plotted against the likelihood of the protein being differentially expressed (−10log values; Y-axes). Grey lines indicate a threshold of S0=0.8. Highlighted are all proteins that are differentially expressed above threshold encoded by Csde1-boundtranscripts (**C**) 2log fold change in average protein expression (iBAQ) following Strap KD versus control Sc002 plotted against 2log fold change in average RNA expression (RPKM) following Strap KD versus control Sc002, for Csde1-bound transcripts only. Red: differential RNA expression at FDR<0.05, blue: differential protein expression at FDR<0.05; green: both RNA and protein differentially expressed. The grey striped line indicates similar change in protein and transcript expression.

In parallel, we analyzed protein expression in Mel cells transduced with lentiviral vectors expressing shRNA against Strap or Sc002 control shRNA. We used label-free quantification (LFQ) of mass spectrometry to compare protein expression in total cell lysates, and analyzed LFQ values with a two-way t-test, which identified 404 proteins as differentially regulated after Strap knockdown with an artificial within groups variance (S0) cutoff of 0.8 (**S6 Table**). Eleven of the differentially regulated proteins after Strap knockdown were encoded by Csde1-bound transcripts (Table 2; Fig. 4B). The lower number of differentially expressed proteins, compared to differentially expressed transcripts, may partly be technical. Proteins expressed at lower levels are not reliably measured, whereas mRNA was measured at greater depth. Yet, the RNA-binding Csde1/Strap complex may differentially express mRNA translation, causing discrepancies between mRNA and protein expression levels. For some differentially expressed proteins, the transcript was not affected by Strap knock down, while in other cases, both transcript and protein expression was affected by Strap knock down.

**Table 2:**
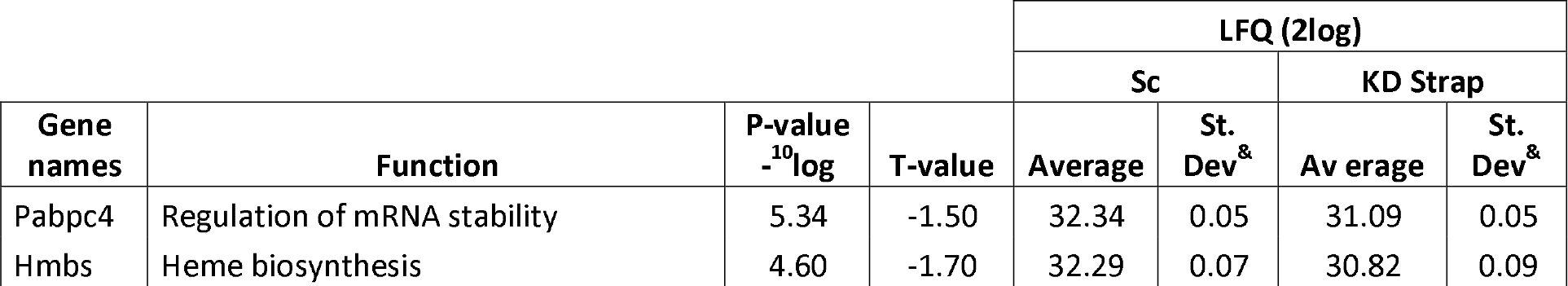
Significant Csde1 target proteins after KD Strap

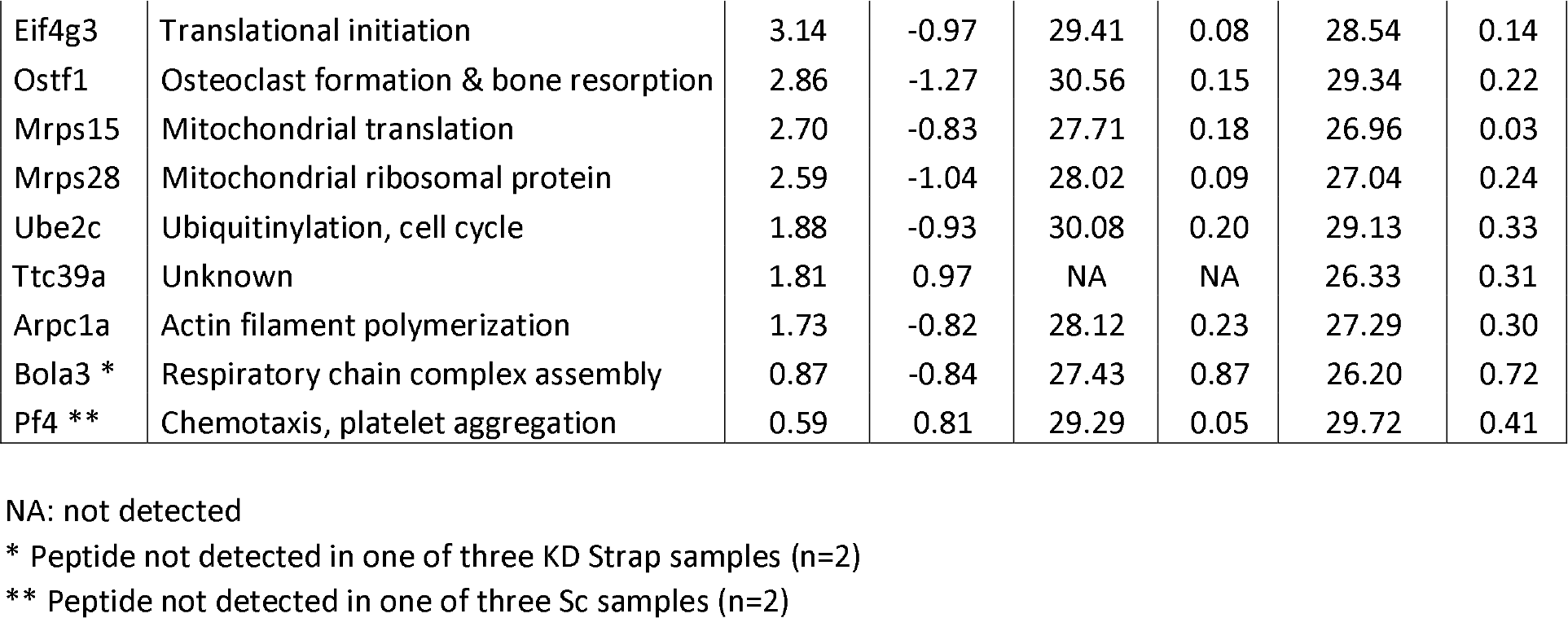

To establish the role of posttranscriptional control, we plotted the fold-change (Strap knock down versus Sc control) in protein (average iBAQ) and transcript (average RPKM) expression for all Csde1-associated transcripts (Fig. 4C). This demonstrated that the Csde1/Strap complex has distinct effects on expression of mRNA and protein of bound transcripts. Expression of mRNA may be reduced, which is compensated by increased protein expression (e.g. Pabpc1) or mRNA expression can be increased whereas protein expression is decreased (e.g. Ptpn7, protein tyrosine phosphatase N7). In both cases, this results in differential mRNA expression, but no statistically significant change in protein expression. Among the downregulated proteins are Pabpc4, a protein that we also found to be associated with Csde1, and that is essential for terminal erythroid differentiation [32]. Pabp proteins directly bind the eIF4G scaffold protein in the eIF4F cap-binding complex to enhance translation [33]. Eif4g3 protein expression is reduced at low Strap levels. It is a variant scaffold protein involved in translation of a selective set of transcripts under hypoxic conditions [34]. Hypoxia-inducible factor-1 (Hif-1)activation regulates at least two other Csde1 targets with reduced protein expression upon Strap KD: heme biosynthesis protein hydroxymethylbilane synthase (Hmbs) and osteoclast stimulating factor 1 (Ostf1) [35,36]. Thus, the Strap/Csde1 complex coordinates mRNA translation and cell cycle divisions, and may also be involved in the hypoxic response in erythroblasts.

### Role of Strap in expression of non-Csde1 associated transcripts

In addition to the regulation of Csde1-associated transcripts, Strap knock down affected the expression of a large number of genes (Fig. 5A). Gene set enrichment analysis (GSEA) on the differentially expressed transcripts upon Strap KD may give an indication of the Csde1 independent role of Strap. For this, we used Genetrail2 [37]. Of particular interest is the number of biological process GO terms related to cell cycle and ribosomal KEGG pathways. It is notable that known Strap pathways such as TGF-B and MAPK were not enriched (**S7 Table**). Because mass spectrometry is less powerful to detect proteins that are expressed at low levels, it is not surprising that we detected fewer differentially expressed proteins. Overrepresentation analysis (ORA) on proteins differentially expressed beyond a threshold of S0=0.8 revealed a diverse array of cellular functions, with biological process GO terms related to cell division, metabolic processes, and vacuole organization of particular prominence within the enriched terms (**S8 Table**). KEGG pathway enrichments include endocytosis, lysosome/peroxisome functionality in addition to several viral-response pathways. The latter can be seen as a predictable consequence to shRNA introduction via viral constructs.

**Fig 5.**
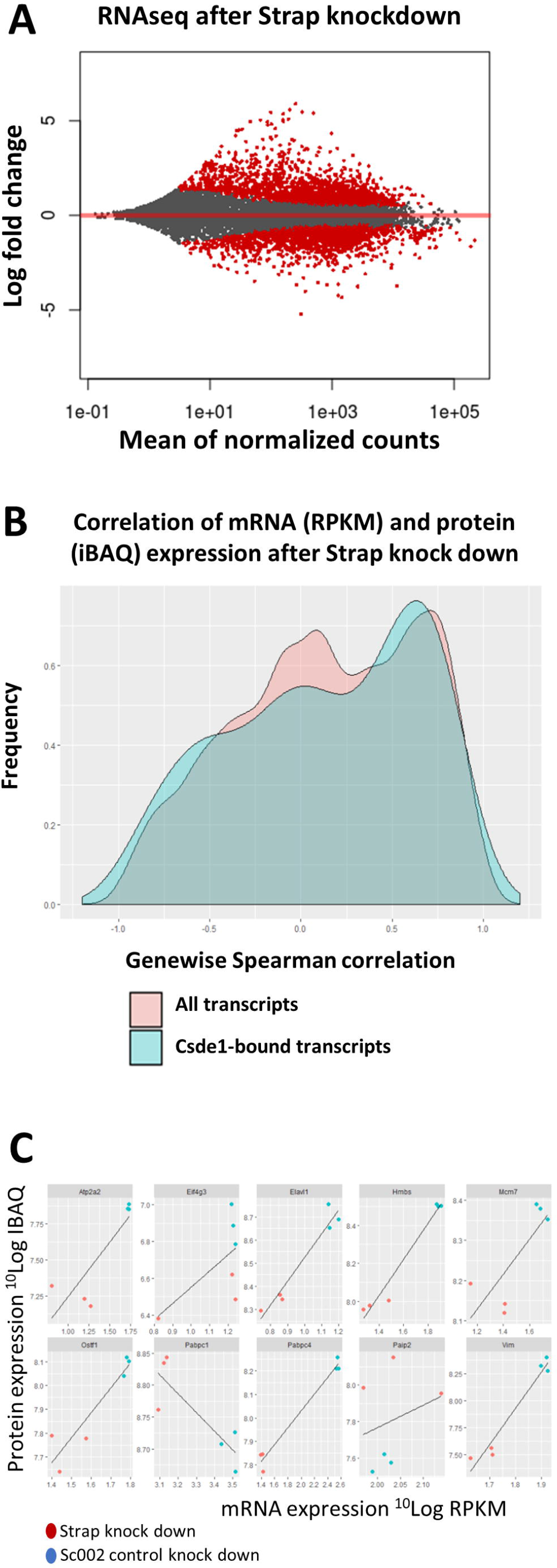
Total transcript and protein expression controlled by Strap. **(A)** MA plot depicting the fold change of mRNA expression (2log cpm) in MEL cells with or without Strap knockdown, plotted against the average expression of the specific mRNA (10log cpm). Transcripts with a FDR<0.05 are highlighted in red (n=3). **(B)** Density plot of genewise Spearman rank correlation coefficients in MEL cells treated with anti-Strap and control shRNA. A comparison of Csde1-bound transcripts (blue) versus all transcripts detectable by both RNAseq and mass spectrometry (red) shows no significant different in the correlation RNA-protein expression. **(C)** Correlation between transcript (10log RPKM) and protein (10log iBAQ) expression levels of select genes.

To assess whether Strap knockdown globally influenced translational regulation, we calculated both sample-wise and gene-wise Spearman correlation coefficients between RNA and protein. Comparisons were limited to genes with at least one valid value within both the RNA sequencing and mass spectrometry datasets. An additional correlation analysis was performed on previously identified Csde1 targets, exclusively. RNA (RPKM) and protein (iBAQ) expression was log10-transformed prior to calculating the correlation coefficient. Strap knock down did not induce significant changes in global correlations in mRNA and protein expression (Table 3). The lack of discernable differences was maintained between all observable genes and previously identified Csde1-targets specifically. A similar lack of differences is evident in a genewise comparison between all observable genes and Csde1 targets (Fig. 5B). Striking differences were observed for gene specific correlations between RNA and protein (**S9 Table**). Pabpc1 protein and RNA expression were negatively correlated with Strap expression in MEL cells expressing anti-Strap or control shRNA, paradoxically indicating a higher protein expression at lower levels of mRNA (Fig. 5C). Interestingly, Strap knockdown reduced mRNA and protein expression of Pabpc4 while increasing expression of its antagonist Paip2, (poly(A) binding protein interacting protein 2). Vim, Atp2a2, Elavi1, Pf4, Ostf1, Hmbs and Mcm7, all of which are implicated in the hypoxic response, are significantly altered in expression at both RNA and the protein level in response to Strap knockdown. By contrast, protein expression of Eif4g3 was strongly reduced by Strap knockdown while RNA expression remains constant, suggesting that Eif4g3 is regulated by Strap at the translational level.

**Table 3:** Sample-wise correlations between RPKM and iBAQ after Strap knockdown

| Sample | Spearman correlation coefficient |  |
| --- | --- | --- |
|  | All genes | Csd1 targets |
| KD Strap a | 0.51 | 0.53 |
| KD Strap b | 0.53 | 0.52 |
| KD Strap c | 0.50 | 0.53 |
| Sc a | 0.51 | 0.53 |
| Sc b | 0.52 | 0.55 |
| Sc c | 0.52 | 0.56 |

## Discussion

The RNA-binding protein Csde1 is essential for erythropoiesis and strongly upregulated in erythroblasts. Reduced expression is associated with DBA [5]. Csde1 binds a subset of transcripts that mainly encode proteins involved in protein homeostasis [15]. The effect of Csde1 on mRNA stability and protein expression of associated transcripts differed. Some were differentially expressed at the transcript level, others at the protein level, which suggested that the nature of the Csde1-containing protein complex is important for the effect of Csde1 on mRNA stability and translation. We identified Strap as a protein that is abundantly associated with Csde1, whereas also Pabpc1 and Pabpc4 are enriched in Csde1 pull downs. Knock down of Strap did not affect Csde1-mRNA interactions, but deregulated mRNA or protein expression of some Csde1-bound transcripts, including Pabpc1 and Pabpc4. This suggests that Pabpc1, and -c4 expression is regulated by the Strap/Csde1 protein complex in a feedback loop. Strap regulated expression of transcripts involved in protein and mRNA homeostasis, and had an effect on the expression of proteins involved in the hypoxia pathway including expression of Eif4g3, a scaffolding component of an alternate eIF4F active under hypoxic conditions, which interacts with Pabpc1 and -c4.

Strap did not affect RNA-binding of Csde1, but it modified RNA and/or protein expression of select Csde1-associated transcripts. At the RNA level, knockdown of Strap altered expression of Csde1 targets involved in translational control, proteasome functionality and cell cycle regulation. The overlap between transcripts involved in cell cycle control and proteasome functionality suggest that Strap may indirectly regulate the cell cycle via post-translational (de)ubiquitination. Strap knock down reduced expression of Ranbp1, which controls the cell cycle via nuclear transport of RNA and proteins [38,39]. Ranbp1 is highly expressed in erythroid progenitors relative to more primitive CD34+ progenitors and displays translation initiation site (TIS) switching in response to eIF2 phosphorylation induced by tunicamycin in erythroid cells [40] Paolini et al., submitted). Strap knockdown also reduced expression of Thoc5, Hmbs, and Ostf1. Thoc5 is a nuclear exporter of mRNAs essential for the maintenance of hematopoiesis [41]. Hmbs catalyzes a crucial step in heme biosynthesis and is oppositely regulated from Hif-1 under hypoxic conditions [35], and Ostf-1 is induced by hypoxia [36,42]. These targets suggest a possible role for Strap in the regulation of terminal erythroid differentiation and the hypoxic response.

Also decreased after Strap knockdown is Vimentin (Vim), which is repressed during terminal erythroid differentiation [43–45]. Biotin pulldown in erythroblasts did not identify Vim as a Csde1-bound transcript [15]. However, an iCLIP approach demonstrated that Csde1 binds the 3′UTR of Vim in human melanoma cells [14]. Knockdown of Csde1 in these cells reduced expression of Vim via decreased ribosome occupancy in melanoma cells. Although the association between Vim and Csde1 is too weak in erythroblasts to be detected, Strap is necessary for enhanced translation of the Vim transcript mediated by Csde1.

Surprisingly, reduction of Strap expression did not result in the altering of Strap-associated KEGG pathways (TGF-B signaling [22,23], MAPK signaling [24], Wnt signaling [25], Notch signaling [26], or SMN complex-mediated splicing [27]). Together, this suggests that the role of Strap in erythropoiesis is mostly in line with Csde1. Csde1 does not sequester Strap away from the nucleus, but the major function of Strap may nevertheless be in the regulation of Csde1 targeted pathways.

Interestingly, we not only substantiated the previously observed interaction of Csde1 with Pabpc1 [11,17], but we also demonstrated interaction of Csde1 with Pabpc4. It is notable that our data did not reproduce other previously identified Csde1 binding partners, such as PTB [7,19], hnRNP C1/C2 [10] or 4E-T [28]. As Csde1 is essential for erythroblast proliferation and differentiation, this suggests that the interaction between Csde1, Strap and Pabpc1/4 is of particular importance during erythropoiesis.

At protein level, loss of Strap resulted in increased Pabpc1 protein expression despite reduction of *Pabpc1* mRNA expression. This finding is similar to what we previously observed for reduced expression of Csde1 itself, and in line with previously published autoregulatory loop in Pabpc1 [15,31]. Loss of Strap also increased the protein:mRNA ratio of Pabpc4, but the strongly reduced expression of *Pabc4* mRNA resulted, nevertheless, in reduced Pabpc4 protein expression. Loss of Strap increased expression of the Pabpc4 antagonist Paip2 (not encoded by a Csde1-bound transcript). Concurrently, loss of Strap reduced expression of elF4g3 expression. Pabpc4, bound to the poly(A) tail, associates with eIF4G, which is part of the pre-initiation scanning complex. This interaction increases mRNA stability and translation [46], and is negatively regulated by competitive binding of Paip2 to eIF4G [47]. Relevantly, eIF4g3 replaces eIF4g1 in an alternate, hypoxic eIF4F complex, which selectively promotes the translation of Hif target transcripts [34]. Of the Pabp family members, Pabpc4 is especially important in erythropoiesis. Pabpc4 stabilizes mRNA transcripts encoding shortened poly(A) tails, including *hα-globin*, *Hif1a*, and *Gata2* [32]. Pabpc4-depleted cells display elevated levels of *c-Kit*, *c-Myb*, *c-Myc*, *CD44*, and *Stat5a*, all genes that are repressed during terminal erythroid differentiation, indicating that Pabpc4 can enhance or inhibit translation in a transcript-specific manner. Given that proteins encoded by Csde1-bound transcripts are generally decreased after Strap knockdown, it is possible is that Csde1-Strap associates with the pre-initiation scanning complex together with Pabpc4 to enhance translation. Taken together with the observed effects of Strap knockdown on Pabpc4 and Paip2, Strap likely amplifies Pabpc4-mediated translational regulation, possibly with a specific role under hypoxic conditions. Alternatively, Csde1-Strap may function as a competitive binding inhibitor, preventing the association of eIF4g3 and Pabpc4 for transcripts normally repressed during erythroid differentiation.

The length of the polyA tail determines how many Pabp molecules can bind to an mRNA transcript. Binding of Pabp protects the polyA tail from deadenylation, for instance by the Cnot1 (CCR4-Not1) deadenylation complex. Cnot1 is downregulated upon Strap knockdown at the protein level [48]. Cnot1 has distinct functions. It has been shown to indirectly interact with both Strap and Csde1 via 4-ET, an eIF4E shuttling protein that transports mRNAs between P-bodies and the cytoplasm [28]. Proteins binding to AU-rich elements in the 3′UTR of transcript can recruit the Cnot1 complex to de-adenylate the polyA tail which may lead to mRNA degradation. The mechanism by which Strap influences the expression of Cnot1 is unclear, but it can be postulated that loss of Strap and subsequent loss of Cnot1 disrupts mRNA degradation and silencing.

Although they are not encoded by Csde1-bound transcripts, Strap influences the expression of several other proteins involved in translational regulation and/or hematopoiesis, the most prominent of which is Elavl1 (HuR), a member of the Elav family of AU-binding protein with well-established roles in hematopoiesis [49]. Strap knock down reduced *Vegfa* expression at the RNA level. Elavl1 binding to AU-rich elements protects transcripts from degradation by preventing the recruitment of Cnot1. For instance, Elavl1 binds to the 3′UTR of Gata1, stabilizing Gata1 translation [50]. Knockdown of Elavla results in the disruption of embryonic erythropoiesis in zebrafish. Elavl1 also stabilizes the *Vegf-a* transcript, a Hif1a-inducible transcript that promotes angiogenesis under hypoxic conditions [51]. Importantly, Elavl1 regulates alternative splicing of *eIF4enif1*, the transcript that encodes 4E-T [52], The absence of Elav1| promotes the expression of the shorter, more stable 4E-T isoform, resulting in drastically increased P-body formation and mRNA turnover of Hif1a, while suppressing angiogenesis via reduced expression of Vegfa. Another member of the Elav family specific to neurons, Elav4 (HuD), is colocalized with the SMN complex in neuronal cells, though this interaction does not depend on the interaction between HuD and RNA [53]. Strap is essential for the assembly of the SMN complex [27], and SMN deficiency is results in the decreased expression of HuD [53]. It is tempting to speculate that knockdown of Strap in erythroblasts results in a comparable loss of HuR via deregulation of the SMN complex.

In conclusion, Strap and Csde1 together form a complex that regulates the translation of transcripts essential for erythropoiesis. Strap does not affect which transcripts are bound by Csde1, but nevertheless alters the translation of select Csde1-bound transcripts involved in the formation of an alternate eIF4F which promotes translation of hypoxic genes. Also regulated by Strap were transcripts associated with terminal erythroid differentiation. Strap may therefore regulate translation during hypoxic erythropoiesis in both a Csde1-dependent or Csde1-independent manner.

## Materials and Methods

### Cell culture

Murine erythroleukemia (MEL) and HEK293T cells were cultured in RPMI, and DMEM respectively (Thermofisher), supplemented with 10% (vol/vol) fetal calf serum (FCS; Bodinco), glutamine and Pen-Strep (Thermofisher). MEL cells expressing BirA, or BirA plus biotag-Csde1 were described previously [5]. Cell number and size were determined using CASY cell counting technology (Roche).

### Lentivirus production and transductions

HEK293Ts were transfected with pLKO.1-puro lentiviral construct containing shRNA sequences for Strap: TRCN0000088837 and a scrambled control shRNA: SHC002 (MISSION TRC-Mm 1.0 shRNA library; Sigma-Aldrich; available on the BloodWeb site), pMD2.G, and pSPAX.2 packaging plasmids (gift of T. van Dijk, Erasmus MC, Rotterdam, The Netherlands) using 0.5M CaCl2 and HEPES (Thermofisher). 72 hours after transduction, viral supernatant was harvested and concentrated using 5% w/v PEG8000 (Sigma). MEL cells were transduced with a multiplicity of infection of 3-5 and addition of 8 μg/mL of Polybrene (Sigma-Aldrich). Transduced cells were selected with 1 μg/ml puromycin 24 hours after transduction.

### Protein-RNA and protein-protein pulldown for Csde1

Biotagged Csde1 containing complexes were collected on streptavidine beads from 10^8^ MEL-BirA or MEL-BirA-Csde1-tag cells (3 biological replicates each) using a previously described protocol [54], with the following modifications. M-270 Dynabeads (Thermofisher; 100μl per 10^8^ cells) were blocked for 1 hour at 4°C in 5% chicken egg albumin and then washed 3× in ice-cold NT2 buffer [50mM Tris-HCI (Sigma-Aldrich), 150mM NaCl (Sigma-Aldrich), 1mM MgCl_2_ (Thermofisher) and 0.05% NP40 (Sigma-Aldrich)]. Cells were lysed in 850μl cold NT2, supplemented by 200U RNAse Out (EMD Bioscience), 400μM vanadyl ribonucleoside complexes (VRC, New England Biolabs) and 20mM EDTA (EM Science), and incubated with the beads for 2 hours at 4°C. Beads were immobilized in a magnet rack, washed 5x with NT2 containing 0.3M NaCl, split into a protein and an RNA fraction. The protein fraction was eluted via boiling in 1x Laemmli buffer (Sigma-Aldrich) for 5 minutes. RNA fractions were purified using Trizol (Invitrogen), precipitated in isopropanol and washed in 75% ethanol.

### SDS-PAGE, Western blotting and silver staining

MEL cells were fractionated into cytoplasmic, nuclear and mitochondrial components using a Cell Fractionation Kit - Standard (ab109719, Abeam), or total MEL cell protein lysates were generated. Proteins were detected via SDS-PAGE and Western blotting as described (Horos et al., 2012). Antibodies used were directed against Strap (sc-136083, Santa Cruz), Csde1 (NBP1-71915, Novus Biological), Stat5 (sc-835, Santa Cruz), Cytochrome C (abll0325, Abeam), Lamin B1 (ab133741, Abeam) and alpha Tubulin (ab4074, Abeam). Fluorescently labeled secondary antibodies for visualization with Odyssey were IRDye 680RD Donkey anti-Rabbit IgG (926-68073, Licor) and IRDye 800CW Donkey anti-Mouse IgG (925-32212, Licor), or using the Pierce enhanced chemiluminescence (ECL) kit (Thermofisher). Silver staining was performed using a SilverQuest™ Silver Staining Kit (LC6070, Thermofisher).

### Mass spectrometry

Eluted peptides were processed as described by [55]. Samples were subjected to mass spectrometry using label-free quantification. All data was analyzed and processed with MaxQuant for peptide identification and quantification [56]. Downstream statistical analysis was performed with Perseus vl.5.1.6 [57]. All peptides matching the reverse database, potential contaminants, and those only identified by site were filtered out. To be considered for analysis, a peptide had to be detectable within all triplicates of at least one clone. Prior to statistical testing, peptide counts were log2 transformed. Because failures to detect a given peptide is sometimes due to insufficient depth, missing values were imputed from the normal distribution with a width of 0.3 and a downshift of 1.8. These values were later de-imputed prior to visualization and production of the final tables. For two-way comparisons between groups, a *t*-test applying an artificial within groups variance of S0=0.8 was used [31]. For all analyses, a Benjamini-Hochberg false discovery rate of < 0.05 was applied. The mass spectrometry proteomics data have been deposited in the ProteomeXchange Consortium via the PRIDE partner repository with the dataset identifier PXD006358 (https://www.ebi.ac.uk/pride/).

### RNA-sequencing

RNA-seq on Csde1-associated transcripts after Strap knockdown was performed by the Leiden Genome Technology Center (LGTC, Leiden), using library preparation following the template-switch protocol (Clontech), and Nextera tagmentation. Samples were split across three MiSeq (lllumina) lanes (2×75bp, paired end). RNA expression by total mRNA sequencing after Strap knockdown was performed by Novogene Co., LTD. on mRNA enriched on oligo(dT) beads. RNA was randomly fragmented, and processed with the NEB Next^®^ Ultra™ RNA Library Prep Kit using random hexamers. The library was sequenced using lllumina HiSeq 2500 (2×150bp, paired end). Sequence quality for both experiments was checked using Fastqc (Babraham Bioinformatics).

Spliced Transcripts Alignment to a Reference (STAR, [58]) was used to align the sequences to the mouse mmlO genomic reference sequence, using the following parameters --outFilterMultimapNmax 20, -- outFilterMismatchNmax 1, --outSAMmultNmax 1, - outSAMtype BAM SortedByCoordinates, quantMode GeneCounts, -outWigType wiggle, -outWigStrand Stranded, --outWigNorm RPM. A gtf file accessed from the UCSC genome browser on 11-Sept-2015 was passed to STAR using -sjdbGTFfile. The read count tables were subjected to differential expression analysis with DESeq2 [59]. DESeq2 implements a negative binomial generalized linear model to identify differential expressed/enriched transcripts. This method normalizes raw counts by adjusting for a size factor to account for discrepancies in sequencing depth between samples. The normalized counts are subsequently subjected to a Wald test with a Benjamini-Hochberg correction for multiple testing (FDR, false discovery rate). When determining whether knockdown of Strap influences which transcripts are bound to Csde1, the following interaction model was applied: ^~^ replicate + shRNA + pulldown + shRNA:pulldown, where shRNA indicates treatment with anti-Strap or control shRNA, replicate indicates the batch, and pulldown indicates the presence or absence of biotagged Csde1. DESeq2 also provides a function for principal component analysis (PCA). Additional visualizations were made using R packages ggplots and pheatmap. Overrepresentation Analysis (ORA) and Gene Set Enrichment Analysis (GSEA) for GO-terms and pathways was performed on significant transcripts with GeneTrail2 [37]. Original sequencing results have been deposited in the BioProject Database under project ID PRJNA379114 (https://www.ncbi.nlm.nih.gov/bioproject/).

### Correlation of RNA and protein expression levels

RNA expression levels were normalized as reads per kilobase of transcript per million mapped reads (RPKM). In mass spectrometry, iBAQ values (as determined via MaxQuant) were normalized via a scaling factor calculated by dividing the sum of intensities from each sample by the intensity sum of a reference sample. A Spearman rank correlation coefficient was calculated between 10log(RPKM) and 10log(iBAQ).

## Acknowledgements

This project was supported by the Landsteiner Foundation for Transfusion Research (LSBR grants 1140 and 1239). We want to thank Drs Henk Buermans and Yavuz Ariyurek, Leiden Genome Technology Centre (LGTC), Leids Universitair Medical Centre (LUMC), for sequencing Csde1 bound transcripts.

## Supporting information captions

**S1 Fig. MA plot on transcripts pulled down by Csde1 in MEL.** The 2log fold change was calculated from average read counts in presence and absence of Strap (cpm, see B) and plotted against 10log average read counts per million (cpm). Highlighted in red are interaction-term significant transcripts. **(A)** MA plot depicting fold change and count abundance of samples containing biotagged Csde1 versus control (BirA, n=3). **(B)** MA plot depicting fold change and count abundance of samples treated with anti-Strap shRNA versus control shRNA (Sc, n=3).

**S2 Fig. Principle component analysis on RNAseq results of Strap knockdown in MEL.** Depicted are both shRNA and replicate groups, indicating that the shRNA is responsible for the majority of variation between samples. PC2 (12%) is the result of minor batch effects.

**S1 Table**: Significant transcripts in Csde1 RIPseq, Csde1-tag vs BirA (control)

**S2 Table**: Significant transcripts in Csde1 RIPseq, Strap knockdown vs control shRNA

**S3 Table**: Interaction-term significant transcripts in Csde1 RIPseq with Strap knockdown

**S4 Table**: Significant transcripts in total mRNA sequencing after Strap knockdown

**S5 Table**: Overrepresentation analysis of Csde1 target transcripts significant after Strap knockdown

**S6 Table**: Significant proteins after KD Strap vs Sc (all proteins)

**S7 Table**: GSEA analysis of Strap knockdown

**S8 Table**: Genetrail2 (ORA) analysis of significant proteins after KD Strap vs Sc (all proteins)

**S9 Table**: Spearman correlation between RPKM and iBAQ after Strap knockdown

